# Rare-Event Sampling using a Reinforcement Learning-Based Weighted Ensemble Method

**DOI:** 10.1101/2024.10.09.617475

**Authors:** Darian T. Yang, Alex M. Goldberg, Lillian T. Chong

## Abstract

Despite the power of path sampling strategies in enabling simulations of rare events, such strategies have not reached their full potential. A common challenge that remains is the identification of a progress coordinate that captures the slow relevant motions of a rare event. Here we have developed a weighted ensemble (WE) path sampling strategy that exploits reinforcement learning to automatically identify an effective progress coordinate among a set of potential coordinates during a simulation. We apply our WE strategy with reinforcement learning to three benchmark systems: (i) an egg carton-shaped toy potential, (ii) an S-shaped toy potential, and (iii) a dimer of the HIV-1 capsid protein (C-terminal domain). To enable rapid testing of the latter system at the atomic level, we employed discrete-state synthetic molecular dynamics trajectories using a generative, fine-grained Markov state model that was based on extensive conventional simulations. Our results demonstrate that using concepts from reinforcement learning with a weighted ensemble of trajectories automatically identifies relevant progress coordinates among multiple candidates at a given time during a simulation. Due to the rigorous weighting of trajectories, the simulations maintain rigorous kinetics.

## Introduction

Path sampling strategies are a powerful class of advanced simulation methods for enhancing the sampling of pathways for rare, barrier-crossing events such as protein (un)folding and (un)binding with rigorous kinetics. ^1^ Prominent examples of path sampling strategies include transition path sampling,^2,3^ transition interface sampling, ^4^ forward flux sampling,^5^ milestoning,^6^ and the weighted ensemble strategy.^7,8^ A major challenge faced by path sampling and many other enhanced sampling methods^9–11^ is the identification of a progress coordinate (also commonly referred to as a reaction coordinate or collective variable), which captures the slowest relevant motion of the barrier-crossing process. ^12^ The process of identifying a progress coordinate often involves a substantial amount of trial-and-error, testing different hypotheses of slow relevant coordinates. Furthermore, the relevant progress coordinate is likely to change at later stages of a multi-step process.

Another class of enhanced sampling methods that requires the identification of progress coordinates is adaptive sampling.^8,11,13,14^ These strategies involve running a large number of trajectories in parallel and replicating (splitting) trajectories at regular short time intervals according to a scoring metric and have been applied in the context of the Folding@home distributing computing project.^15^ In contrast to path sampling strategies, however, trajectories generated by adaptive sampling do not directly yield rigorous rates^13^ and must be combined with the construction of either Markov state models (MSMs)^14,16–18^ or generalized master-equation-based models^19,20^ to estimate rates and other long-timescale observables such as state populations. Adaptive sampling strategies can be broadly classified into Markov state model (MSM)-based and machine learning (ML)-based methods. MSM-based methods such as least-count adaptive sampling are useful for the exploration of configurational space.^21^ ML-based methods such as reinforcement learning (RL)-based methods are useful for balancing the exploration of configurational space with exploitation of pathways towards a target state and have been used to identify effective progress coordinates among multiple candidates on-the-fly while running the simulation.^13^ Examples of RL-based adaptive sampling methods include FAST,^22^ AdaptiveBandit,^23^ TSLC,^24^ and REAP.^25^

Inspired by adaptive sampling efforts, we present here a weighted ensemble (WE) path sampling strategy that employs reinforcement learning (WE-RL) to automatically identify an effective progress coordinate during a simulation. Rather than applying ML to learn the progress coordinate from the atomic coordinates of sampled conformations,^12,26–28^ our WE-RL method employs the strategy of periodically identifying the most effective coordinate among multiple proposed candidates during a simulation.^29–31^ WE path sampling is a “splitting strategy” like adaptive sampling where promising trajectories are replicated at regular short time intervals. In contrast to adaptive sampling, WE sampling involves the rigorous assignment of statistical weights to the trajectories to ensure that no bias is introduced into the dynamics, enabling direct calculations of rates from the simulations themselves. While the progress coordinate for WE simulations is typically divided into bins, our WE-RL method involves a “binless” framework in which conformations are automatically clustered by similarity along the progress coordinate at periodic intervals, circumventing the need to manually position bins.

To demonstrate the power of our RL-based WE method, we focus on three benchmark systems: (i) an egg carton-shaped toy potential to test exploration of unknown regions, (ii) an S-shaped toy potential to test the generation of pathways toward a target state, and (iii) a dimer of the HIV-1 capsid protein (C-terminal domain) to test conformational sampling of an atomistic system. To enable rapid testing of the latter, we ran discrete-state synthetic molecular dynamics trajectories that were propagated using a generative, fine-grained Markov state model based on extensive conventional MD simulations.

## Theory

In this section, we outline key features of the weighted ensemble (WE) strategy and then present an adaptation of this approach that utilizes reinforcement learning (WE-RL) to sample underexplored regions of phase space. As a step towards developing our WE-RL method, we first adapted the WE strategy to utilize least-count sampling (WE-LC), another feature besides reinforcement learning that was inspired by efforts involving adaptive sampling of free energy landscapes.^21,32,33^

### The Weighted Ensemble Strategy

The weighted ensemble (WE) strategy involves running a large number of weighted trajectories in parallel and iteratively applying a resampling procedure to efficiently generate region-to-region transitions toward a target state. ^7,8,34^ Typically, regions are defined as bins along a progress coordinate that is intended to capture the slowest relevant motion of the process of interest. The resampling procedure is applied at short fixed time intervals *τ* with the goal of evenly distributing trajectories along the progress coordinate by maintaining an equal number of trajectories per bin. To achieve this goal, the procedure replicates (splitting) trajectories that transition to a less-visited bin and occasionally terminates (merging) trajectories that occupy a more frequently-visited bin. Importantly, trajectory weights are rigorously tracked such that no bias is introduced into the dynamics, enabling direct calculations of rates.

The methods below exploit a key feature of the WE strategy: trajectory weights are independent of the progress coordinate such that the progress coordinate (and bin positions) can be switched on-the-fly during a simulation. While the original WE strategy involves the use of bins along a progress coordinate, the strategy can also be adapted for a “binless” framework^35^ in which clusters of conformations instead of bins are used to guide the WE resampling procedure. Below, we present the use of a binless framework for a least-count WE sampling (WE-LC) method and a reinforcement learning-based weighted ensemble (WE-RL) method. For both of these methods, the number of generated clusters is tunable. If no input is provided, the number of clusters will be determined using a heuristic function.^24^

### Least-Count Sampling with Weighted Ensembles of Trajectories

Our least-count weighted ensemble sampling method (WE-LC) involves periodically applying a clustering procedure such as *k*-means during a simulation, splitting trajectories in lower-count clusters and merging trajectories in higher-count clusters. We outline the steps of the WE-LC workflow below.

1. Initiate multiple weighted trajectories.
2. Run dynamics of the ensemble in parallel for a fixed time interval *τ*.
3. Cluster across all candidate progress coordinates of the trajectories in the current WE iteration using *k*-means clustering, generating the cluster set *C*, where each individual cluster *c* ∈ *C*.
4. Sort the resulting clusters based on member counts.
5. Split within the low count cluster(s) and merge within the high count cluster(s), maintaining a constant number of trajectories per WE iteration.
6. Repeat steps 2-5 for *N* WE iterations.

### Reinforcement Learning with Weighted Ensembles of Trajectories

Our reinforcement learning-based WE strategy (WE-RL) involves the periodic application of reinforcement learning^25^ to identify the most effective progress coordinate among multiple candidates at a given time during a simulation. From RL, we use the concept of a sampling “policy” *π*, which maps an agent’s “state” (*S*) within the environment to an “action” (*A*): *π* : *S* → *A*. The RL definition of state should not be confused with the biophysical definition referring to protein conformational states. In our implementation, we seek to identify a sampling policy which is dependent on a corresponding set of *K* progress coordinates (*π*_*K*_, *K* = {*θ*_1_, *θ*_2_, …, *θ*_*k*_}), where the agent is the ensemble of weighted trajectories, the state is the current set of progress coordinate data available for the current WE iteration, and the action is the trajectory resampling procedure.

Our WE-RL workflow involves steps 1-4 in the WE-LC workflow with the following additional steps.

5. Select a subset of the lowest count clusters where *C*_*LC*_ ⊂ *C*. Selecting a subset of low count clusters ensures that the worst this algorithm performs should still be comparable to least-count adaptive sampling.^21^
6. Set the weight *w*_*i*_ for each *θ*_*i*_ ∈ *K*, where *w*_*i*_ ∈ [0, 1]. Weights for each progress coordinate can be set ahead of time, and otherwise will default to 1*/k*, where *k* is the total number of progress coordinates for the sampling policy *π*_*K*_.
7. Given the set of *K* progress coordinates for policy *π*_*K*_, calculate the reward for each cluster, noting that *c*_*j*_ ∈ *C*_*LC*_.

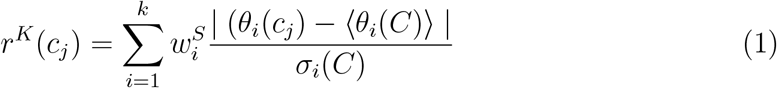

Where 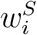 represents the weight of each progress coordinate for the state space *S, θ*_*i*_(*c*_*j*_) is the progress coordinate calculated for cluster *c*_*j*_, ⟨*θ*_*i*_(*C*)⟩ is the arithmetic mean of *θ*_*i*_ for all *c* ∈ *C*, and *σ*_*i*_(*C*) represents the standard deviation of *θ*_*i*_ for all *c* ∈ *C*.

8. We then calculate the cumulative reward:

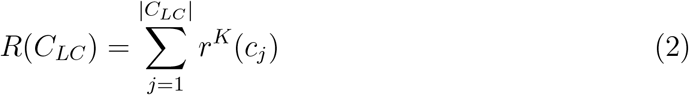

Where the sum is over each element in the least-count cluster set (*C*_*LC*_).
9. Equation 2 is maximized using the Sequential Least SQuares Programming (SLSQP) method in SciPy^36^ to optimize the contribution of each progress coordinate weight 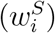. The following conditions were enforced as constraints during the optimization process: ∑*w*_*i*_ = 1 and 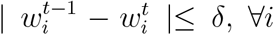, where 0 *< δ <* 1 and *t* represents the current WE iteration while *t* − 1 represents the previous WE iteration. In short, the progress coordinate weights always sum to 1 and the *δ* parameter prevents the progress coordinate weights from changing drastically. Using the updated weights, step 7 is repeated to calculate the updated reward values of each cluster.
10. We then split the trajectories within clusters with the highest reward, while merging the trajectories in the highest count cluster(s), maintaining a constant number of trajectories per WE iteration.
11. Repeat steps 2-10 for *N* WE iterations.

## Methods

### Weighted Ensemble Simulations

All WE simulations were carried out using the open-source and highly scalable Weighted Ensemble Simulation Toolkit with Parallelization and Analysis (WESTPA) 2.0 software package.^37^ WE simulations were performed in a non-equilibrium steady-state ensemble, where trajectories that reached the target state were “recycled” back to the initial state while maintaining the same trajectory weight. WE data analysis and plotting was done using the WEDAP package.^38^ Based on the Hill relation,^39,40^ rate constants were calculated from the probability flux into the target state by tracking the total weight among successful (recycled) trajectories per unit time.

### Conventional Simulations

To assess the advantage of periodically applying the WE resampling procedure, a comparable number of trajectories were run in parallel using conventional simulations (without resampling) as a reference set. For example, for a custom binless WE simulation with 80 trajectories, the reference set consisted of 80 conventional trajectories.

### An Egg Carton-shaped Toy Potential

The egg carton-shaped potential consists of a series of low-energy wells separated by modest energy barriers (Figure 1.A). The potential in one dimension is defined as:

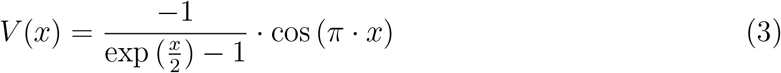

**Figure 1:**
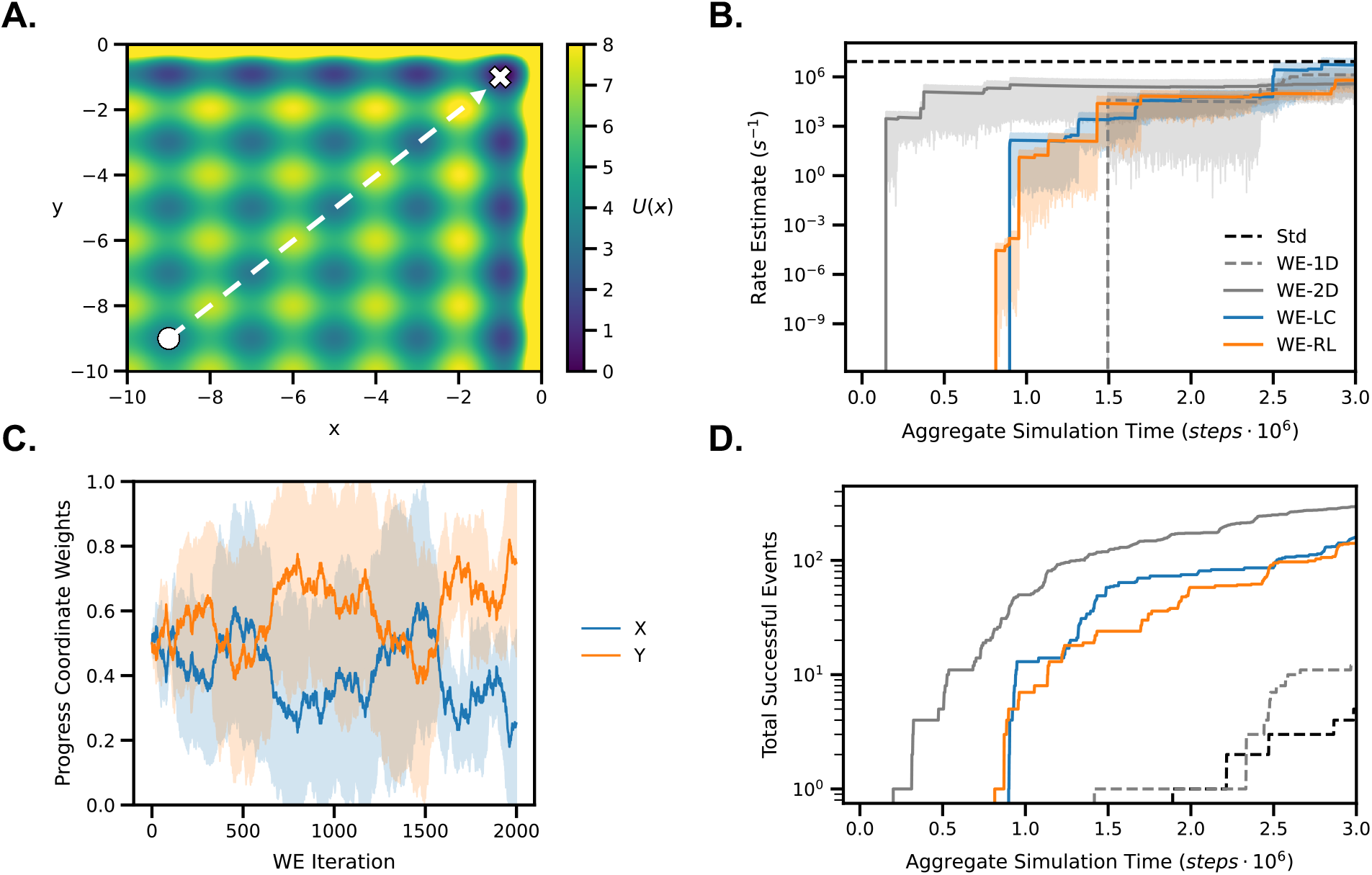
The performance of various WE simulation schemes along an egg carton-like toy potential. (A) The egg carton-shaped potential energy function from a two-dimensional version of Equation 3. The initial and target states are marked by a white circle and a white X, respectively. (B) Rate estimates of each WE simulation scheme, compared to the conventional, parallel dynamics reference rate (dashed, horizontal black line). Uncertainties for each scheme were calculated using Bayesian bootstrapping^43^ and 95% credibility regions are shown in translucent coloration. (C) Evolution of progress coordinate weights during the WE-RL simulations, where the average (solid lines) and one standard deviation (translucent fill) are shown. (D) The amount of successfully recycled trajectories over time for each sampling scheme. The identities of each line are the same as depicted in the legend on panel (B).

This potential was used with both the X and Y positions to create a two-dimensional egg carton-shaped potential, and dynamics were propagated using the overdamped Langevin equation:

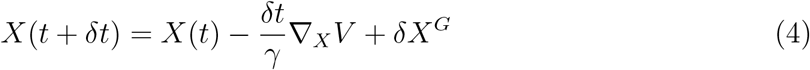

where *γ* is the friction coefficient, *δt* is the time step, and *δX*^*G*^ is a random displacement with zero mean and variance 2*γk*_*B*_*Tδt* with *δt* = 5 · 10^*−*5^ and reduced units of *γ* = 1 and *k*_*B*_*T* = 1. Reflecting boundaries were placed at -10 and 0 in both the X and Y dimensions and WE simulations were run with a dynamics propagation interval (*τ*) of 20 integrator steps per iteration. All simulations were initialized with X and Y positions at -9.5 and recycling was carried out once a trajectory reached the target state past -1.5 in both X and Y positions.

For the one-dimensional rectilinear WE binning scheme, 4 bins were positioned along the Y dimension at -7, -5, and -3 with a target of 20 trajectories per bin. For the two-dimensional WE binning scheme, 24 bins were symmetrically positioned along both the X and Y positions at -7, -5, -3, and -1.5, with a target of 4 trajectories per bin. For the reference set of conventional simulations and the binless WE methods, 80 trajectories were maintained throughout the simulation. Each WE simulation method was run for 5 independent trials. The conventional simulations were run for a length equivalent to 5000 WE iterations to ensure convergence in the estimated rate constant. The WE-1D, WE-2D, WE-LC, and WE-RL simulations were each run for 2000 WE iterations.

### An S-shaped Toy Potential

The S-shaped toy potential consists of a pathway that “snakes” along small-to-medium sized barriers and meta-stable intermediate states (Figure 2.A), defined as:

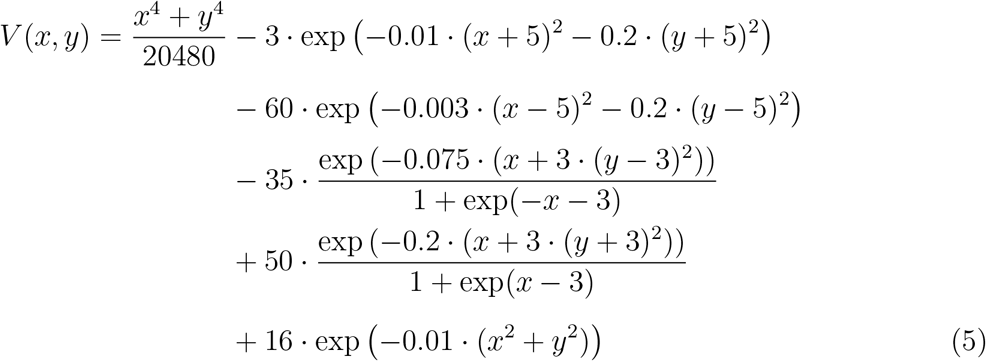

**Figure 2:**
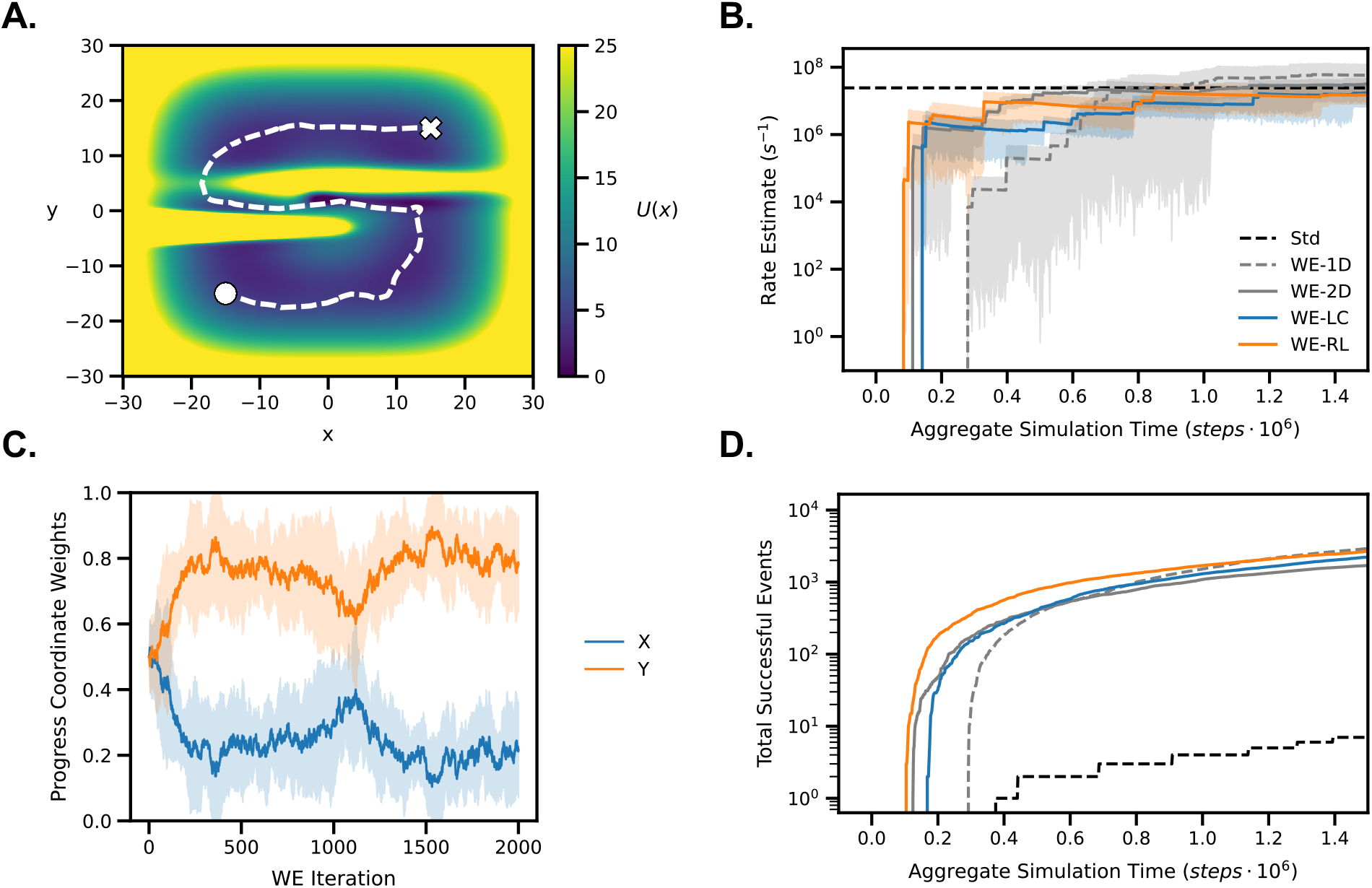
The performance of various WE simulation schemes for simulating an S-shaped pathway. (A) The S-shaped potential energy function (Equation 5). The initial and target states are marked by a white circle and a white X, respectively. (B) Rate estimates of each WE simulation scheme, compared to the conventional, parallel dynamics reference rate (dashed, horizontal black line). Uncertainties for each scheme were calculated using Bayesian bootstrapping^43^ and 95% credibility regions are shown in translucent coloration.(C) Evolution of progress coordinate weights during the WE-RL simulations, where the average (solid lines) and one standard deviation (translucent fill) are shown. (D) The amount of successfully recycled trajectories over time for each sampling scheme. The identities of each line are the same as depicted in the legend on panel (B).

Dynamics propagation was carried out using the OpenMM dynamics engine, ^41^ where each individual simulation consisted of a single particle with a mass of 1 AMU. Each simulation was run at a temperature of 300 K with a friction coefficient of 1 ps^-1^ using a Langevin integrator. Simulations were initialized at X and Y positions of -15. All WE simulations used a resampling time interval (*τ*) of 20 integrator steps. To reach a non-equilibrium steady state, trajectories were recycled back to the initial state positions after reaching X and Y values greater than 10.

For both the binless WE methods tested and the corresponding reference set of conventional simulations, the total number of trajectories was maintained at 80. As another point of reference, WE simulations were carried out using two different binning approaches. The first approach used a one-dimensional binning scheme (WE-1D) with 10 bins placed along the X position at intervals of 5 from -20 to 20 with a target of 8 trajectories per bin. The second approach involved a two-dimensional binning scheme (WE-2D) where 6 bins were positioned along both the X and Y positions at intervals of 10 from -20 to 20, yielding a total of 36 bins and a target count of 4 trajectories per bin.

For each resampling scheme, five independent WE simulations were run. The reference set of conventional simulations was run to a length that was equivalent to 10000 WE iterations to ensure convergence of the reference rate constant. The WE-1D, WE-2D, WE-LC, and WE-RL simulations were each run for 2000 WE iterations.

### Synthetic Molecular Dynamics of a HIV-1 Capsid Protein Dimer

A fine-grained Markov state model was constructed using ∼90 µs of conventional MD simulations of the HIV-1 capsid protein C-terminal domain dimer, a 100-ps MSM lag time *τ*, and a non-standard, stratified clustering scheme to better preserve kinetic properties at shorter lag times.^42^ Discrete-state synthetic molecular dynamics trajectories were propagated along a Markov chain using the Markov state model, as implemented in the Synthetic Dynamics (SynD)^42^ Python package. At each *τ*, each microbin was back-mapped to representative structure of that microbin in a multi-dimensional progress-coordinate space, which was defined using the following six features: (i) orientation angle 1, (ii) orientation angle 2, (iii) T188-T188 distance, (iv) C_2_ angle, (v) the W184 *χ*_1_ angle of monomer 1, and (vi) the W184 *χ*_1_ angle of monomer 2. Simulations began at the correspondingly ordered progress coordinate values of (i) 39.8°, (ii) 42.7°, (ii) 15.3 Å, (iv) 70.6°, (v) 173°, and (vi) 176°. Trajectories were recycled upon reaching the target state, which was defined by both orientation angles being less than 10°, the T188-T188 distance being less than 5 Å, the C_2_ angle being greater than 40°, and the W184 *χ*_1_ angles being between -95° and -40°. Trajectories were propagated 1 step for each WE iteration, which corresponds to the MSM lag time of 100 ps.

For our binned WE simulation scheme, 10 bins were placed along the T188-T188 distance in 1 Å intervals from 5 to 14 Å with a target of 8 trajectories in each bin. The WE-LC and WE-RL methods were run with 80 trajectories per iteration and the following four progress coordinates: orientation angle 1, orientation angle 2, T188-T188 distance, and the C_2_ angle. Min-max scaling of input progress coordinates was carried out to ensure that different ranges of input progress coordinates were clustered with equal contributions. Overall, 5 independent WE simulations using a binned WE approach, the WE-LC method, and the WE-RL method were run for 2,000 WE iterations each (∼0.24 ms of aggregate “synthetic” simulation time). The reference rate constant estimate was obtained directly from the MSM transition matrix, where the mean first passage time was 3.83 µs.

## Results and Discussion

We first examined the performance of our modified WE methods with two toy potentials: (i) an egg carton-like potential with small barriers and wells symmetrically distributed along the X and Y dimensions (Figure 1.A) to test the efficiency of general phase space exploration, and (ii) a narrowing, S-shaped potential where regions outside of the “S” shape are out of reach (Figure 2.A), thus testing the ability of our WE algorithms to find and exploit a single pathway to a target state. Finally, we explore a more realistic molecular system by sampling alternate dimer orientations of the HIV-1 capsid protein C-terminal domain (CTD) using synthetic MD trajectories (Figure 3.A).

**Figure 3:**
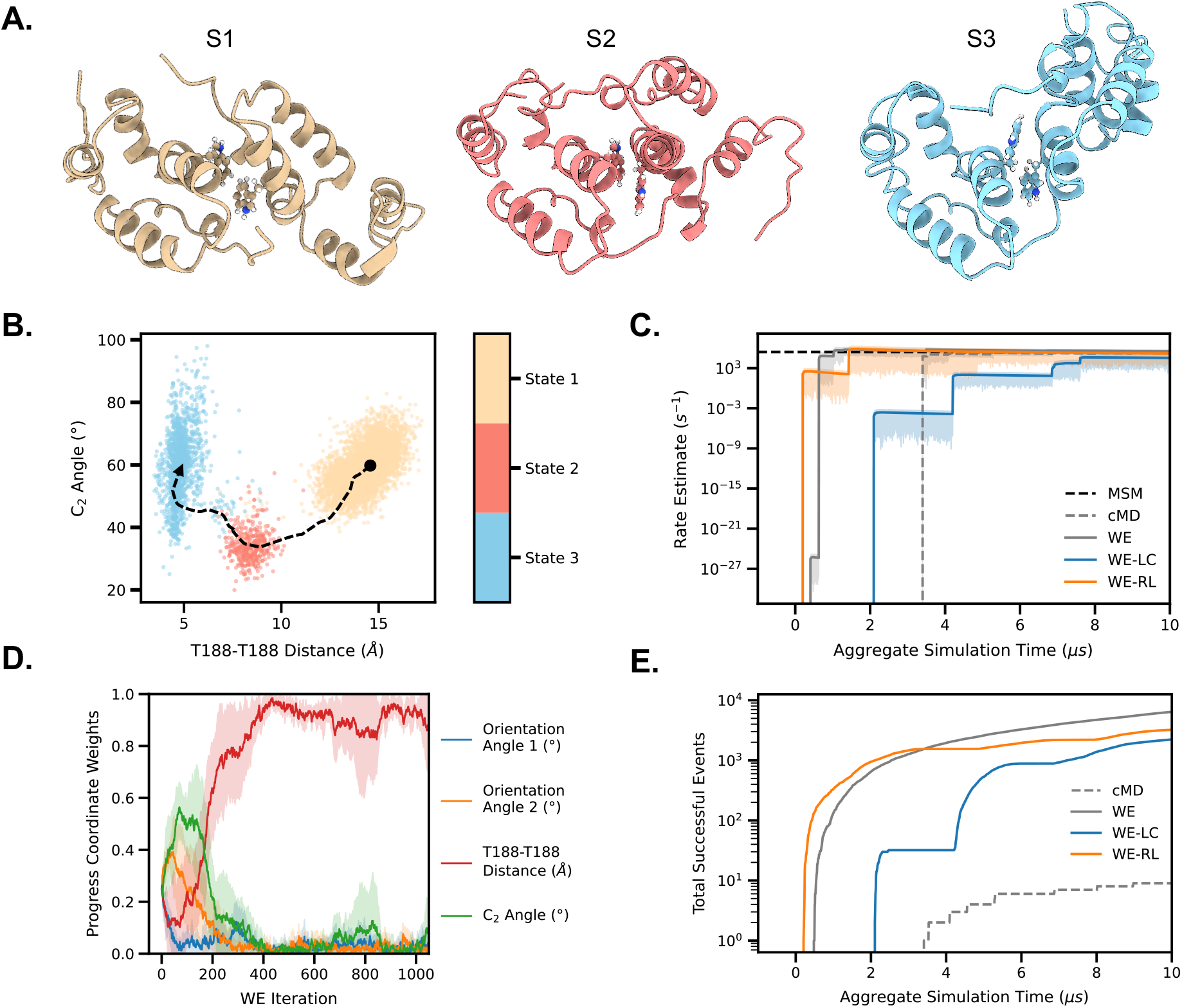
The performance of various WE simulation schemes for conformational sampling of the HIV-1 capsid protein CTD dimer. (A) Representative states of the capsid protein CTD dimer. (B) The pathway of interconversion between states 1, 2, and 3 along two candidate progress coordinates. (C) Rate estimates of each simulation scheme, compared to the reference rate from the MSM (dashed, horizontal black line). Uncertainties for each scheme were calculated using Bayesian bootstrapping^43^ and 95% credibility regions are shown in translucent coloration. (D) Early evolution of progress coordinate weights during the WE-RL simulations, where the average (solid lines) and one standard deviation (translucent fill) are shown for all four progress coordinates. (E) The amount of successfully recycled trajectories over time for each sampling scheme.

### Exploration: Simulations using an Egg Carton-shaped Potential

Using an egg carton-shaped potential (Figure 1.A), we compared WE simulations using bins in either one or two dimensions with our binless WE-LC and WE-RL methods. For the former, we binned in only one dimension to represent a “bad” choice of progress coordinate (neglecting movements in the second dimension), while the two-dimensional binning scheme is ideal for exploring the entire potential. Typically, not all degrees of freedom in a simulation system are known *a priori*. Therefore, in the case of the two-dimensional binning example, we anticipated it would perform more efficiently than our binless WE methods. Indeed, both WE-LC and WE-RL outperformed WE simulations using the one-dimensional binning scheme. However, they under-performed compared to WE simulations employing the ideal two-dimensional binning scheme, specifically in terms of yielding a converged rate estimate that was comparable to the reference rate from extensive conventional simulations (Figure and the amount of successful events (Figure 1.D).

For the goal of exploration, WE-LC and WE-RL performed similarly, likely due to the purely exploration-based sampling process, where adaptive sampling has previously been shown to be highly efficient.^21,32,33^ For the egg carton potential, the WE-RL method maintains equal importance contributions to both the X and Y progress coordinates (Figure 1.C), which is logical considering the symmetry of the potential. Thus, as expected, the general task of symmetric phase-space exploration did not benefit from the exploration/exploitation balance of the WE-RL method.

### Exploitation: Simulations using an S-shaped Potential

Using a narrowing S-shaped potential (Figure 2.A), we compared the ability of binned WE simulations with our binless WE methods for exploiting the S-shaped pathway. We again use a one-dimensional binning scheme to represent a “bad” choice of progress coordinates and a two-dimensional binning scheme to present a “good” choice of progress coordinates. We find that the one-dimensional binning scheme performs poorly as expected, the WE-LC and WE simulations with two-dimensional binning perform similarly in rate constant convergence (Figure 2.B) and number of successful events (Figure 2.D), and the WE-RL simulations are able to perform on par with the two-dimensional binning scheme simulations in terms of rate constant convergence, while being able to sample more early transition events.

For the task of path exploitation, WE-RL performed better than WE-LC and the binned WE simulation schemes. WE-RL simulations were shown to place more importance on the Y dimension of the progress coordinate over time (Figure 2.C), which may correlate to the presence of the highest barrier being in the Y dimension (Figure 2.A). When sampling along a single pathway, the balance of exploration and exploitation provided by the WE-RL algorithm may have an advantage compared to the other WE simulation schemes.

### Conformational Dynamics of the HIV-1 Capsid Protein Dimer

The HIV-1 capsid protein (CA) self-assembles into the mature viral capsid, which is crucial for successful host cell infection and encapsulates the viral RNA genome within a flexible hexameric lattice, enclosed at the ends of the ovoid by pentameric subunits.^44,45^ The flexibility of the capsid is attributable to the CA, which connects adjacent hexameric and pentameric subunits along the inside of the core through dimerization of the CA-CTD. We previously explored the multi-state ensemble of the CA-CTD dimer (Figure 3.A) through ^19^F NMR experiments and an extensive set of both WE and conventional MD simulations (unpublished), identifying a set of progress coordinates to describe the flexibility of the dimer interface. The ∼90 µs of WE simulation-seeded conventional MD simulations from our previous study were used to build an MSM of the CA-CTD dimer with discrete transitions between multiple dimer states. Using this MSM, we generated “synthetic” trajectories^42^ of the CA-CTD dimer, to test our modified WE algorithms by sampling the transition from state 1 to state 3 (Figure 3.B).

We compared the ability of WE-LC and WE-RL to sample the CA-CTD dimer interface using multiple candidate progress coordinates against a binned WE simulation with a single progress coordinate. The WE-LC and WE-RL simulations used a four-dimensional progress coordinate consisting of orientation angles 1 and 2, the T188-T188 distance, and the C_2_ angle, while the binned WE simulation used just the T188-T188 distance, which we have previously shown to be a “good” progress coordinate that describes the entire ensemble of dimer states (unpublished). We found that while WE-LC simulations captured events faster than conventional MD (cMD), both binned WE and WE-RL simulations recorded more transition events (Figure 3.C) and achieved faster convergence of rate constants to the reference MSM rate (Figure 3.E) compared to both cMD and WE-LC. Notably, WE-RL simulations sampled transition events slightly faster than even the binned WE approach, but required marginally more simulation time for rate constant convergence. The performance of our binless WE approaches in sampling the CA-CTD dimer conformations mirrors the trends observed in our simulations of the S-shaped potential, likely due to the path-like nature of the multi-state CA-CTD conformational transition network.

Using the WE-RL method, we consistently found that T188-T188 distance was the most important progress coordinate to focus sampling along, while also identifying the C_2_ angle as a useful early coordinate (Figure 3.D). The initial focus of the WE-RL simulations on C_2_ angle is consistent with the state distribution of this transition process (Figure 3.B) and may explain the early jump in successful events since the C_2_ angle is known to promote the CA-CTD state 1 to state 2 transition. The T188-T188 distance, however, can effectively capture the entire transition process from S1 to S3, which is reflected in the overall progress coordinate weight distribution as the WE-RL simulations progressed. In the CA-CTD dimer system, WE-RL is able to identify ideal early and overall progress coordinate contributions.

## Conclusions

We have developed a WE path sampling strategy that integrates features of adaptive sampling (WE-LC) and reinforcement learning (WE-RL) to automate the selection of progress coordinates. In a binless framework, reinforcement learning was used to “learn” the importance of each candidate progress coordinate during a WE simulation. Unlike adaptive sampling and reinforcement learning algorithms that do not employ weighted trajectories,^13,22–25^ our WE strategies enable the direct calculation of rate constants. By testing the exploration and exploitation capabilities of our modified WE algorithms on model potentials, we found that WE-LC and WE-RL could outperform the binned WE strategy, especially when effective progress coordinates and bin placements were unknown. For the S-shaped potential used to assess path exploitation, the WE-RL algorithm slightly outperformed the two-dimensional binning scheme, which fully captured the potential energy landscape. This improvement may be due to the WE-RL algorithm “learning” to prioritize the Y dimension of the progress coordinate, where the largest barrier was present. In simulations of the HIV-1 capsid protein CTD dimer, WE-RL performed comparably to WE simulations using a “good” progress coordinate and was able to identify this same coordinate as the most critical. Overall, WE-RL is a promising initial approach when system-specific information is unknown and can be used to identify ideal progress coordinates for subsequent simulations.

## Notes

LTC serves on the scientific advisory board of OpenEye Scientific Software.

## Acknowledgement

We thank all members of the Chong lab for valuable discussions.

## References

(1) Chong, L. T.; Saglam, A. S.; Zuckerman, D. M. Path-sampling strategies for simulating rare events in biomolecular systems. 43, 88–94.

(2) Dellago, C.; Bolhuis, P. G.; Csajka, F. S.; Chandler, D. Transition path sampling and the calculation of rate constants. The Journal of Chemical Physics 1998, 108, 1964–1977.

(3) Bolhuis, P. G.; Chandler, D.; Dellago, C.; Geissler, P. L. TRANSITION PATH SAMPLING: Throwing Ropes Over Rough Mountain Passes, in the Dark. Annual Review of Physical Chemistry 2002, 53, 291–318.

(4) van Erp, T. S.; Moroni, D.; Bolhuis, P. G. A novel path sampling method for the calculation of rate constants. The Journal of Chemical Physics 2003, 118, 7762–7774.

(5) Allen, R. J.; Valeriani, C.; Wolde, P. R. t. Forward flux sampling for rare event simulations. J. Phys.: Condens. Matter 2009, 21, 463102.

(6) Faradjian, A. K.; Elber, R. Computing time scales from reaction coordinates by milestoning. The Journal of Chemical Physics 2004, 120, 10880–10889.

(7) Huber, G. A.; Kim, S. Weighted-ensemble Brownian dynamics simulations for protein association reactions. Biophysical Journal 1996, 70, 97–110.

(8) Zuckerman, D. M.; Chong, L. T. Weighted Ensemble Simulation: Review of Methodology, Applications, and Software. Annual Review of Biophysics 2017, 46, 43–57.

(9) Torrie, G. M.; Valleau, J. P. Nonphysical sampling distributions in Monte Carlo freeenergy estimation: Umbrella sampling. 23, 187–199.

(10) Laio, A.; Parrinello, M. Escaping free-energy minima. 99, 12562–12566.

(11) Hénin, J.; Lelièvre, T.; Shirts, M. R.; Valsson, O.; Delemotte, L. Enhanced Sampling Methods for Molecular Dynamics Simulations [Article v1.0]. 4, 1583.

(12) Fu, H.; Bian, H.; Shao, X.; Cai, W. Collective Variable-Based Enhanced Sampling: From Human Learning to Machine Learning. The Journal of Physical Chemistry Letters 2024, 15, 1774–1783.

(13) Kleiman, D. E.; Nadeem, H.; Shukla, D. Adaptive Sampling Methods for Molecular Dynamics in the Era of Machine Learning. J. Phys. Chem. B 2023, 127, 10669–10681.

(14) Husic, B. E.; Pande, V. S. Markov State Models: From an Art to a Science. Journal of the American Chemical Society 2018, 140, 2386–2396.

(15) Pande, V. S.; Baker, I.; Chapman, J.; Elmer, S. P.; Khaliq, S.; Larson, S. M.; Rhee, Y. M.; Shirts, M. R.; Snow, C. D.; Sorin, E. J. et al. Atomistic protein folding simulations on the submillisecond time scale using worldwide distributed computing. Biopolymers 2003, 68, 91–109.

(16) Pande, V. S.; Beauchamp, K.; Bowman, G. R. Everything you wanted to know about Markov State Models but were afraid to ask. Methods 2010, 52, 99–105.

(17) Suárez, E.; Wiewiora, R. P.; Wehmeyer, C.; Noé, F.; Chodera, J. D.; Zuckerman, D. M. What Markov State Models Can and Cannot Do: Correlation versus Path-Based Observables in Protein-Folding Models. Journal of Chemical Theory and Computation 2021, 17, 3119–3133.

(18) Prinz, J.-H.; Wu, H.; Sarich, M.; Keller, B.; Senne, M.; Held, M.; Chodera, J. D.; Schütte, C.; Noé, F. Markov models of molecular kinetics: Generation and validation. The Journal of Chemical Physics 2011, 134, 174105.

(19) Cao, S.; Montoya-Castillo, A.; Wang, W.; Markland, T. E.; Huang, X. On the advantages of exploiting memory in Markov state models for biomolecular dynamics. The Journal of Chemical Physics 2020, 153, 014105.

(20) Dominic, A. J.; Sayer, T.; Cao, S.; Markland, T. E.; Huang, X.; Montoya-Castillo, A. Building insightful, memory-enriched models to capture long-time biochemical processes from short-time simulations. Proceedings of the National Academy of Sciences 2023, 120, e2221048120.

(21) Weber, J. K.; Pande, V. S. Characterization and Rapid Sampling of Protein Folding Markov State Model Topologies. J. Chem. Theory Comput. 2011, 7, 3405–3411.

(22) Zimmerman, M. I.; Bowman, G. R. FAST Conformational Searches by Balancing Exploration/Exploitation Trade-Offs. J. Chem. Theory Comput. 2015, 11, 5747–5757.

(23) Pérez, A.; Herrera-Nieto, P.; Doerr, S.; De Fabritiis, G. AdaptiveBandit: A Multiarmed Bandit Framework for Adaptive Sampling in Molecular Simulations. J. Chem. Theory Comput. 2020, 16, 4685–4693.

(24) Buenfil, J.; Koelle, S. J.; Meila, M. Tangent Space Least Adaptive Clustering. ICML 2021 Workshop on Unsupervised Reinforcement Learning. 2021.

(25) Shamsi, Z.; Cheng, K. J.; Shukla, D. Reinforcement Learning Based Adaptive Sampling: REAPing Rewards by Exploring Protein Conformational Landscapes. J. Phys. Chem. B 2018, 122, 8386–8395.

(26) Lee, H.; Ma, H.; Turilli, M.; Bhowmik, D.; Jha, S.; Ramanathan, A. DeepDriveMD: Deep-Learning Driven Adaptive Molecular Simulations for Protein Folding. https://arxiv.org/abs/1909.07817.

(27) Tian, H.; Jiang, X.; Xiao, S.; La Force, H.; Larson, E. C.; Tao, P. LAST: Latent Space-Assisted Adaptive Sampling for Protein Trajectories. Journal of Chemical Information and Modeling 2023, 63, 67–75.

(28) Leung, J. M. G.; Frazee, N. C.; Brace, A.; Bogetti, A. T.; Ramanathan, A.; Chong, L. T. Unsupervised learning of progress coordinates during weighted ensemble simulations: Application to millisecond protein folding. bioRxiv 2024,

(29) Tiwary, P.; Berne, B. J. Spectral gap optimization of order parameters for sampling complex molecular systems. 113, 2839–2844.

(30) Ribeiro, J. a. M. L.; Bravo, P.; Wang, Y.; Tiwary, P. Reweighted autoencoded variational Bayes for enhanced sampling (RAVE). 149, 072301.

(31) Mehdi, S.; Smith, Z.; Herron, L.; Zou, Z.; Tiwary, P. Enhanced Sampling with Machine Learning. 75, 347–370.

(32) Singhal, N.; Snow, C. D.; Pande, V. S. Using path sampling to build better Markovian state models: Predicting the folding rate and mechanism of a tryptophan zipper beta hairpin. The Journal of Chemical Physics 2004, 121, 415–425.

(33) Bowman, G. R.; Ensign, D. L.; Pande, V. S. Enhanced Modeling via Network Theory: Adaptive Sampling of Markov State Models. J. Chem. Theory Comput. 2010, 6, 787–794.

(34) Zhang, B. W.; Jasnow, D.; Zuckerman, D. M. The “weighted ensemble” path sampling method is statistically exact for a broad class of stochastic processes and binning procedures. 132, 054107.

(35) Donyapour, N.; Roussey, N. M.; Dickson, A. REVO: Resampling of ensembles by variation optimization. 150, 244112.

(36) Virtanen, P.; Gommers, R.; Oliphant, T. E.; Haberland, M.; Reddy, T.; Cournapeau, D.; Burovski, E.; Peterson, P.; Weckesser, W.; Bright, J. et al. SciPy 1.0: fundamental algorithms for scientific computing in Python. Nature Methods 2020, 17, 261–272.

(37) Russo, J. D.; Zhang, S.; Leung, J. M. G.; Bogetti, A. T.; Thompson, J. P.; DeGrave, A. J.; Torrillo, P. A.; Pratt, A. J.; Wong, K. F.; Xia, J. et al. WESTPA 2.0: High-Performance Upgrades for Weighted Ensemble Simulations and Analysis of Longer-Timescale Applications. Journal of Chemical Theory and Computation 2022, acs.jctc.1c01154.

(38) Yang, D. T.; Chong, L. T. WEDAP: A Python Package for Streamlined Plotting of Molecular Simulation Data. Journal of Chemical Information and Modeling 2024, 64, 5749–5755.

(39) Hill, T. Free Energy Transduction and Biochemical Cycle Kinetics; Dover Publications, 2004.

(40) Bhatt, D.; Zuckerman, D. M. Beyond Microscopic Reversibility: Are Observable Nonequilibrium Processes Precisely Reversible? J. Chem. Theory Comput. 2011, 7, 2520–2527.

(41) Eastman, P.; Swails, J.; Chodera, J. D.; McGibbon, R. T.; Zhao, Y.; Beauchamp, K. A.; Wang, L. P.; Simmonett, A. C.; Harrigan, M. P.; Stern, C. D. et al. OpenMM 7: Rapid development of high performance algorithms for molecular dynamics. PLOS Computational Biology 2017, 13, e1005659.

(42) Russo, J. D.; Zuckerman, D. M. Simple synthetic molecular dynamics for efficient trajectory generation. https://arxiv.org/abs/2204.04343.

(43) Mostofian, B.; Zuckerman, D. M. Statistical Uncertainty Analysis for Small-Sample, High Log-Variance Data: Cautions for Bootstrapping and Bayesian Bootstrapping. Journal of Chemical Theory and Computation 2019, 15, 3499–3509.

(44) Campbell, E. M.; Hope, T. J. HIV-1 capsid: the multifaceted key player in HIV-1 infection. 13, 471–483.

(45) Mattei, S.; Glass, B.; Hagen, W. J. H.; Kräusslich, H.-G.; Briggs, J. A. G. The structure and flexibility of conical HIV-1 capsids determined within intact virions. 354, 1434–1437.

